# Transcriptional signatures underlying divergent lifestyles of endophytic and pathogenic fungi in early colonisation of wheat roots

**DOI:** 10.64898/2026.03.13.711015

**Authors:** Stella Moren-Rosado, Rowena Hill, Tania Chancellor, Rachel Rusholme Pilcher, Neil Hall, Kim E. Hammond-Kosack, Mark McMullan

## Abstract

Wheat take-all is a root disease which devastates crop yields, caused by the ascomycete fungus *Gaeumannomyces tritici*. The closely related root endophyte, *G. hyphopodioides*, has been found to induce local host defence responses which confer protection against take-all and reduce disease severity. Chancellor et al. (2024) investigated host transcriptional response to early colonisation by each of these two fungi. Using this RNA-seq dataset in conjunction with newly available *Gaeumannomyces* reference genomes, we have completed the picture by characterising the fungal transcriptional activity underpinning these different lifestyles. Even at early time points, their transcriptional profiles differ: *G. hyphopodioides* shows signs of transcriptional reprogramming between 4 and 5 days post inoculation (dpi), mirroring the wheat response, whereas *G. tritici* expression varied very little between these two time points despite progressing into the vasculature, instead exhibiting a stealthy expression profile dominated by gene downregulation at earlier time points. Moreover, GO term enrichment in this study identified a stress-response unique to *G. hyphopodioides*, which may explain the formation of its subepidermal vesicles (SEVs), putative resting structures that are a key difference between the pathogen and non-pathogen, alongside upregulation of many putative effectors and CAZymes. The enrichment of a key lignin-degrading CAZyme may contribute to the lack of stress-response identified in *G. tritici*, allowing fungal hyphae to overcome localised host lignification. These findings highlight the transcriptional basis of colonisation differences and are a step towards understanding how closely related fungi with different lifestyles modulate their interactions within a common host and tissue.

## Introduction

In the face of climate change, crop security is threatened by plateauing yields, strained natural resources, soil health degradation, and emerging crop diseases (Mc Carthy et al. 2018; Fones et al. 2020). The annual crop yield loss to plant diseases is estimated to be up to 30% globally (Savary et al. 2019). Pathogens affecting root health are expected to thrive best in areas where winter conditions are becoming wetter and milder, likely to further reduce crop yields (Palma-Guerrero et al. 2021). Plants must constantly interact with a complex and dynamic soil microbiome in the rhizosphere where even closely related fungi can adopt diverse lifestyles, ranging between pathogenic, non-pathogenic endophytes, and beneficial symbionts. The ability to navigate these rhizosphere dynamics successfully is known to be a major influence on plant health and development (Raaijmakers et al. 2009). Soil microbial communities are home to the greatest reservoir of biological diversity known so far in the world (Berendsen et al. 2012) and are gaining traction in crop health and pathogen control research, as we strive to reduce our dependence on chemical pest control and fertilisers. General disease suppression is linked to the total microbial biomass in soil (Weller et al. 2002), so understanding the interactions between all actors will be a crucial avenue of research for enhancing crop security without compromising biodiversity.

The soil-borne ascomycete fungus, *Gaeumannomyces tritici* (*Magnaporthaceae*) is the pathogenic agent of wheat take-all, which is considered the most important wheat root disease worldwide (Freeman and Ward 2004). Its runner hyphae spread among hosts producing multiple infections along the root surface, where hyphopodia then penetrate the root epidermis. Hyphae colonise the root cortex and finally invade the central stele, containing the plant vascular tissue. The pathogenic *G. tritici* infection forms characteristic necrotic lesions, and the subsequent destruction of the vasculature severely compromises water and nutrient uptake (Palma-Guerrero et al. 2021).

Crop rotations which include non-cereal species are currently the predominant measure used to reduce *G. tritici* inoculum levels in the soil (Palma-Guerrero et al. 2021). Fungicide seed treatments are available and while efficacy is limited to the seedling phase this is enough to assist root development and improve yield. However, naturally occurring fungicide resistance has been reported in up to 30% of *G. tritici* strains (Freeman et al. 2005; Yun et al. 2012). Efforts to mine the rhizosphere for potential microbial biocontrol agents may offer less costly and more ecologically sensitive approaches towards crop resilience. Beneficial microorganisms can provide protection by either direct antagonism of the pathogen or by inducing host resistance mechanisms. For example, *Bacillus subtilis* strains have been found to lower take-all disease incidence equal to, or more than Silthiofam fungicide (Yang et al. 2018, 2015a). Production of antimicrobial compounds by *Pseudomonas* spp. has been well-characterised and has been linked to spontaneously induced soil suppressiveness, resulting in take-all decline (TAD) (Freeman and Ward 2004; Kwak and Weller 2013; Mehrabi et al. 2016). Non-pathogenic fungal species *Slopeiomyces cylindrosporus* and *Gaeumannomyces hyphopodioides*, which are closely related to *G. tritici*, are also promising take-all biocontrol candidates (Chancellor 2022; Speakman and Lewis 1978). *G. hyphopodioides* has been found to reduce take-all disease levels and *G. tritici* biomass by inducing local resistance in the host, when it colonises the root first (Chancellor et al. 2024; Speakman and Lewis 1978).

*G. hyphopodioides* is an endophyte which also colonises wheat roots but differs from *G. tritici* in that its colonisation is restricted to the root cortex (Fig. 1). It is not possible at this stage to attribute the cause of this confinement to either the host or the fungus (or both). However, *G. hyphopodioides* does not go on to infect the vascular tissue, does not cause disease symptoms and, as such, it is described as “non-pathogenic”. Another novelty of the *G. hyphopodioides* colonisation in comparison to *G. tritici* is the production of melanised, swollen fungal cells known as subepidermal vesicles (SEVs) in the root cortex (Chancellor et al. 2024). Often referred to as “growth cessation-structures”, SEVs are thought to be formed where hyphal growth is arrested, perhaps as a stress response, although it is not clear if this is due to locally induced wheat defences or nutrient deficiency (Freeman and Ward 2004; Chancellor et al. 2024). It is not typically addressed in the literature, but *G. hyphopodioides* could be considered a member of the polyphyletic guild of root-dwelling fungi known as dark septate endophytes (DSEs), as these melanised SEVs are likely synonymous with the characteristic microsclerotia of DSEs (Malicka et al. 2022; Chancellor 2022). Some DSEs can engage cooperatively with the plant host despite not having specialised nutrient exchange structures, and as such are hypothesised to represent a phase on the evolutionary path towards mycorrhizal symbiosis (Ruotsalainen et al. 2022).

**Figure 1.**
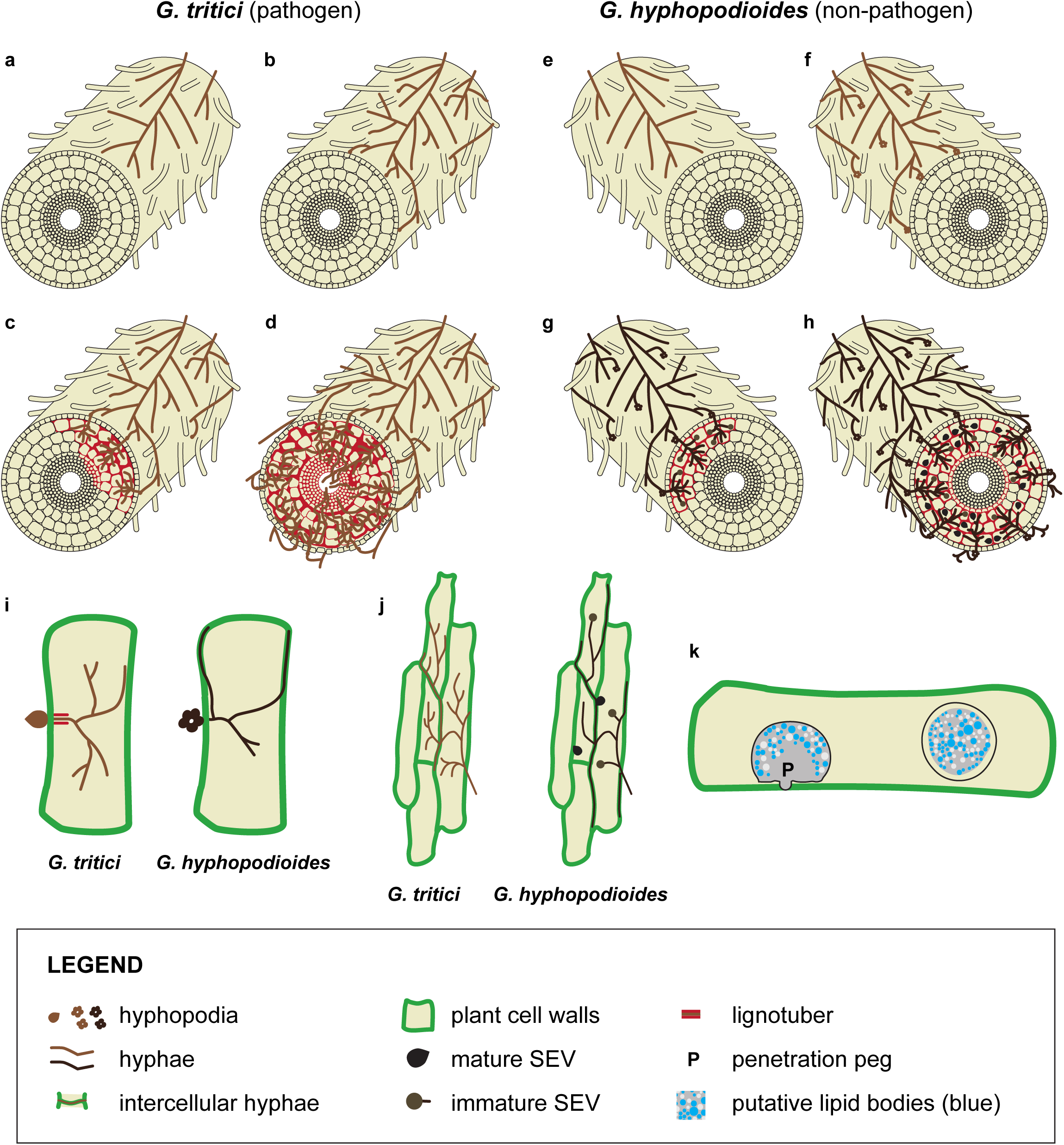
Contrasting root invasion strategies of pathogenic and endophytic *Gaeumannomyces* species. Four stages of *G. tritici* **(a-d)** and *G. hyphopodioides* **(e-h)** invasion show initial similarities followed by lifestyle divergence. Both pathogen and endophyte runner hyphae grow on root surfaces **(a, e)** and penetrate the epidermis via hyphopodia - simple for *G. tritici* **(b)** or multi-lobed for *G. hyphopodioides* **(f)**. Both colonise the cortex inter- and intracellularly **(c, g)** but diverge at the endodermis: *G. tritici* penetrates through to destroy vascular tissue despite lignotuber formation **(c-d)**, while *G. hyphopodioides* remains confined to the cortex, developing pigmented hyphae and subepidermal vesicles (SEVs; **g-h**). **(i-j)** Intracellular invasion patterns differ between pathogen (left) and endophyte (right), with *G. tritici* causing severe disorganisation and lignotubers, while *G. hyphopodioides* grows along cell peripheries forming SEVs with penetration pegs. (k) Endophytic SEVs contain putative lipid bodies.

To investigate the mechanism of this take-all control, Chancellor et al. (2024) completed the first comparative analysis of the host transcriptional response to colonisation by both *G. tritici* and *G. hyphopodioides*. Extensive transcriptional reprogramming of the wheat host associated with triggering of a localised immune response was observed in tissue colonised with the non-pathogen, *G. hyphopodioides*, which was not seen in response to the pathogen, *G. tritici*. This expression change occurred in parallel to the formation of *G. hyphopodioides* SEVs in the root cortex (Chancellor et al. 2024). At the time of the previous study, transcriptomic profiling of *G. tritici* and *G. hyphopodioides* themselves during colonisation was not possible, due to the lack of high-quality reference fungal genomes.

The present study aims to expand the scope of this valuable expression dataset which contains the fungal transcriptomes in addition to that of the host. Analysing fungal reads using the newly available *Gaeumannomyces* reference genomes of the same isolates (Hill et al. 2025) provides an exciting opportunity for comparative molecular exploration of the lifestyle and colonisation differences between pathogenic *G. tritici* and endophytic *G. hyphopodioides*. Leveraging this dataset offers insights into numerous unanswered questions: Namely, are early colonisation processes conserved in both species? Is *G. hyphopodioides* colonisation confined to the cortex by the host’s or its own biology? Is *G. tritici* able to suppress the host immune response or do hyphae simply evade detection? These are important next steps in developing our molecular understanding of the take-all system’s interspecific interactions, involving host, pathogen, and non-pathogenic endophyte, which offers a unique microcosm to study fungal lifestyle differences, transience, and the potential of microbial biocontrol measures.

## Materials and Methods

### RNA-seq experimental design and methods

RNA-seq was performed by Chancellor et al. (2024), in brief: host wheat plants of cv. Chinese Spring were precision inoculated with fungal plugs of either *G. tritici* (isolate Gt17LH(4)19d1 = Gt-19d1) or *G. hyphopodioides* (isolate NZ.129.2C.17 = Gh-2C17) in an agar plate system, with 5 replicates for each treatment (Table 1). Root pieces at the same fungal colonisation stage were snap frozen for RNA extraction at three different time points: 2-, 4-, and 5-days post inoculation (dpi). *In vitro* fungal samples of *G. tritici* and *G. hyphopodioides* were prepared in Potato Dextrose Broth (PDB) with 5 replicates of each for RNA extraction, which provided a control for the fungal read mapping in this study – these were performed as part of the original investigation (Chancellor et al. 2024) but are published here for the first time (Table 1). mRNA library preparation was carried out by Novagene and sequenced by Illumina NovaSeq. Please refer to Chancellor et al. (2024) for further details on experimental set up. Control fungal RNA-seq data introduced here are deposited in the NCBI Gene Expression Omnibus (GEO) under accession GSE324493. The published dual RNA-seq data can be found under accession GSE242417.

**Table 1.** Summary of treatments for RNA-seq.

| Treatment | Time point(s) | Replicates | Data reference |
| --- | --- | --- | --- |
| Wheat - | 2, 4, 5 dpi | A, B, C, D, E | (Chancellor et al. 2024) |
| Wheat + Gh-2C17 | 2, 4, 5 dpi | A, B, C, D, E | (Chancellor et al. 2024) |
| Wheat + Gt-19d1 | 2, 4, 5 dpi | A, B, C, D, E | (Chancellor et al. 2024) |
| PDB + Gh-2C17 | 4 days growth | A, B, C, D, E | This study |
| PDB + Gt-19d1 | 4 days growth | A, B, C, D, E | This study |

### RNA-seq Read Mapping and Quantification

The raw sequencing data fastq files from Chancellor et al. (2024) were downloaded from the Sequence Read Archive (SRA) and combined with the previously unpublished control data for the two fungi (Table 1). Read data quality was evaluated using FastQC v1.19 (Andrews 2018). The quality of base calls was extremely high, with none falling into the poor-quality range. Reads were left untrimmed to avoid the biases in gene expression quantification introduced by trimming (Liao and Shi 2020; Williams et al. 2016). The reference genomes for wheat cv. Chinese Spring (IWGSC RefSeqv2.1; Zhu et al. 2021), Gt-19d1 and Gh-2C17 (Hill et al. 2025) were indexed and RNA-seq reads were aligned to the corresponding reference genomes using HISATv2.1.0 (Kim et al. 2019). Alignment statistics were summarised using Samtools flagstat v1.18 (Danecek et al. 2021). Transcript abundance was quantified using StringTie v2.1.1 (Kovaka et al. 2019) and the raw gene count matrix was constructed using prepDE.py script (available at: https://github.com/gpertea/stringtie/blob/master/prepDE.py).

### Exploratory Data Quality Checks

To determine the minimum fungal read count threshold for meaningful differential gene expression analysis, the proportion of total reads aligning to the fungal reference genomes at each time point was compared with the level of mis-mapping between uninoculated host samples and the fungal references. Dobon et al. (2016) excluded time points (1, 2, & 3 dpi) because the percentage of reads mapping to the fungal reference genome (up to 1%) was similar to the proportion that mis-mapped in the uninoculated control. In the present study, however, the proportion of mis-mapping reads at the early 2 dpi time point was 100-fold lower (< 0.01% for most samples; Supplementary Fig. S1). Additionally, histograms of average transcripts per million (TPM) at each time point were plotted to ensure the transcriptome was represented comprehensively within the reads, without systematic bias (Supplementary Fig. S2). While we advocate some caution in interpreting the data at 2 dpi due to overall low read counts, we felt these quality checks justified retaining all samples in downstream analyses.

### Differential Expression Analysis

All downstream analyses were performed using R v4.1.2 (R Core Team 2024). The raw gene count matrix was filtered to contain only genes with at least 10 counts in at least 3 samples. The differential gene expression of both *G. tritici* and *G. hyphopodioides* at each time point, compared to their corresponding controls, was analysed. The R package DESeq2 v1.40.2 (Love et al. 2014) was used for normalisation and calling of differentially expressed genes, the results of which were filtered for log2 fold change of > 2 or < -2, together with a Benjamini-Hochberg adjusted p-value < 0.05 (Benjamini and Hochberg 1995) implemented by DESeq2 to control for multiple testing. The full table of significant DEGs can be found in the Supplementary Datasheet. Variance-stabilised principle component analyses (PCAs) of the top 500 most variable genes were generated using the vst and plotPCA functions from DESeq2 for each of the two species separately, as well as a combined PCA using shared single copy orthogroups previously inferred by Hill et al. (2025).

### GO Term and Gene Set Enrichment Analysis

Gene Ontology (GO) term enrichment analysis of significantly up- and down-regulated fungal genes at each time point was performed in R, using the package topGO v2.52.0 (Alexa and Rahnenfuhrer 2022). GO terms were retrieved from the functional annotations (Hill et al. 2025) and Fisher’s exact test was used to determine significantly enriched GO terms. Gene set enrichment analysis (GSEA) of carbohydrate-active enzymes (CAZymes; *G. hyphopodioides* n=224, *G. tritici* n=211), candidate secreted effector proteins (CSEPs; *G. hyphopodioides* n=139, *G. tritici* n=126) and biosynthetic gene clusters (BGCs; *G. hyphopodioides* n=4, *G. tritici* n=5) was performed using the fgsea v1.26.0 package (Korotkevich et al. 2016) with genes ranked according to the ‘stat’ column containing the Wald statistic from the DESeq2 results. All data visualisation was done in RStudio using the ggplot2 v4.0.0 package (Wickham 2016).

## Results

### Sequencing depth increases over time as fungal colonisation proceeds

Transcript data from wheat roots colonised with either the pathogen *G. tritici* or the endophyte *G. hyphopodioides* were originally used to understand host response to infection (Chancellor et al. 2024). Here, we first assessed the feasibility of this dual host-fungus data to understand the fungal side of the interaction. At the earliest sequenced infection stage, 2 days post inoculation (dpi), both the proportion of total reads attributed to the fungi (< 0.5%; Fig. 2a) and the average read counts (∼300-600k; Fig. 2b) were low for both *G. tritici* and *G. hyphopodioides*. By 4 dpi, the number and proportion of reads increased for both *G. tritici* and *G. hyphopodioides*, congruent with the fungal colonisation progression seen in the corresponding host root microscopy (Chancellor et al. 2024). Strikingly, at 5 dpi there was a dramatic increase in the number of reads mapping to *G. hyphopodioides,* accounting for nearly a quarter of the total (Fig. 2a), with one of the replicates contributing a remarkably substantial 40.7% of reads. This sudden increase was not seen in the pathogenic *G. tritici*, where both read count and proportion remained relatively consistent, with only a small increase from 2.8% to 4.0% of the total reads between 4 and 5 dpi.

**Figure 2.**
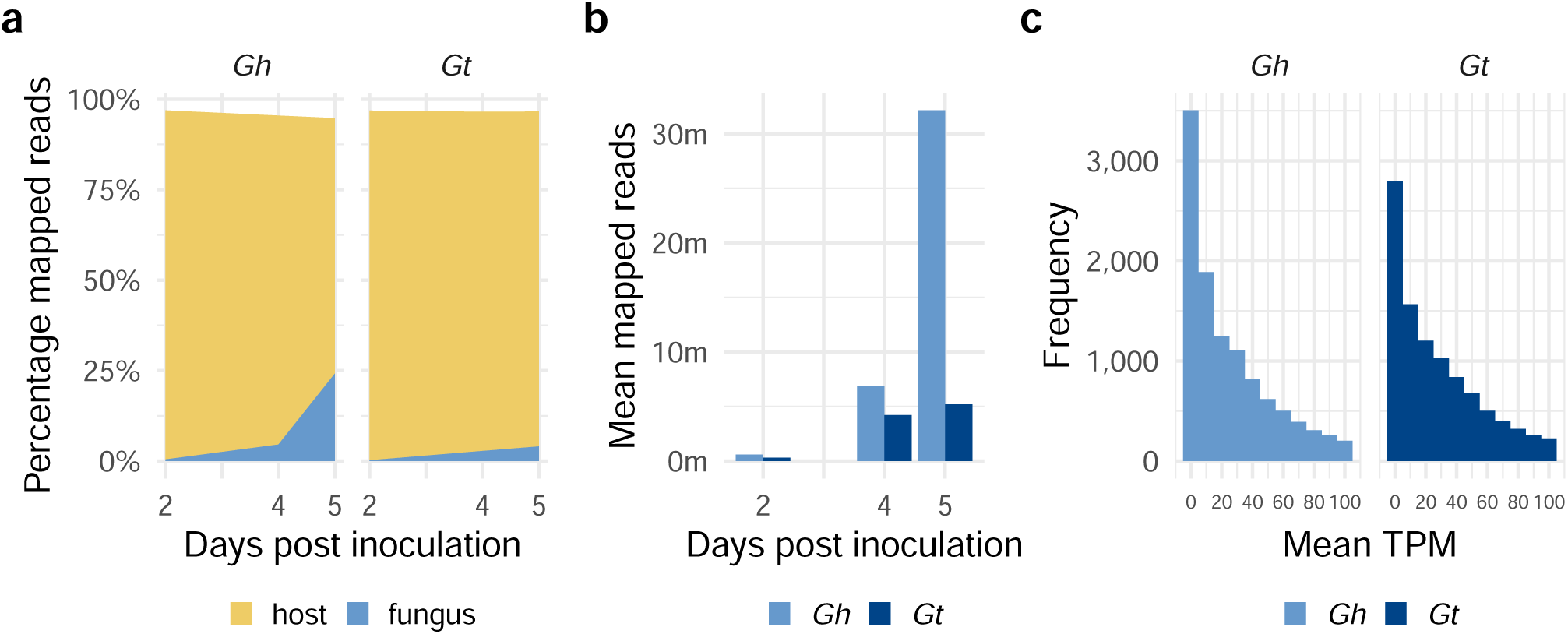
**(a)** Percentage of the total RNA reads mapping to the wheat host versus either the non-pathogenic *G. hyphopodioides* (*Gh*) or pathogenic *G. tritici* (*Gt*) fungus at each time point, averaged across replicates. **(b)** Mean number of reads mapped to each fungus at each point. **(c)** Histograms of the mean transcripts per million (TPM) per gene for each fungus (see Supplementary Fig. S2 for separate histograms for each time point).

### Fungal transcriptomes were well-represented within the RNA-seq reads

For meaningful differential gene expression analysis, it is important to ensure transcriptomes are properly represented. Histograms of average transcripts per million (TPM) at each time point show an optimal distribution of transcript abundance and gene expression for both fungi, without systematic bias towards a small number of highly expressed genes (Fig. 2c). This optimal distribution for both fungi includes 2 dpi where read counts were lower. The vast majority of genes were either lowly- or not- expressed, which aligns with DESeq2’s assumption that most genes will remain unchanged between conditions (Love et al. 2014). Nevertheless, there were still sufficient genes with higher expression levels, facilitating informative differential gene expression analysis and ensuring that DESeq2’s estimations of dispersion and fold changes would be appropriate for this dataset.

### Diverging colonisation strategies between pathogen and non-pathogen

Differential gene expression analysis was completed by comparing the fungi at each stage of colonisation with their respective control samples grown in PDB media. We used principal component analysis (PCA) to assess expression divergence for both the pathogenic *G. tritici* and non-pathogenic *G. hyphopodioides* across the time course of early colonisation. Both fungi had clear separation of expression profiles for the early 2 dpi time point, excepting one sample for *G. hyphopodioides* that clustered with the 4 dpi samples (Fig. 3a). Interestingly, this sample had been flagged as an outlier in the original host transcriptome analysis based on a higher number of reads mapping to the *G. tritici* genome relative to other replicates at this time point (Chancellor et al. 2024). However, for *G. hyphopodioides*, 4 and 5 dpi each formed distinct clusters, showing a marked transcriptional shift across the period when lifestyle diverges and SEVs form in *G. hyphopodioides*, while 4 and 5 dpi were clustered together for *G. tritici*. When visualised together, using shared single copy orthogroup genes, the variation in expression was accounted for by species (Supplementary Fig. S3b).

**Figure 3.**
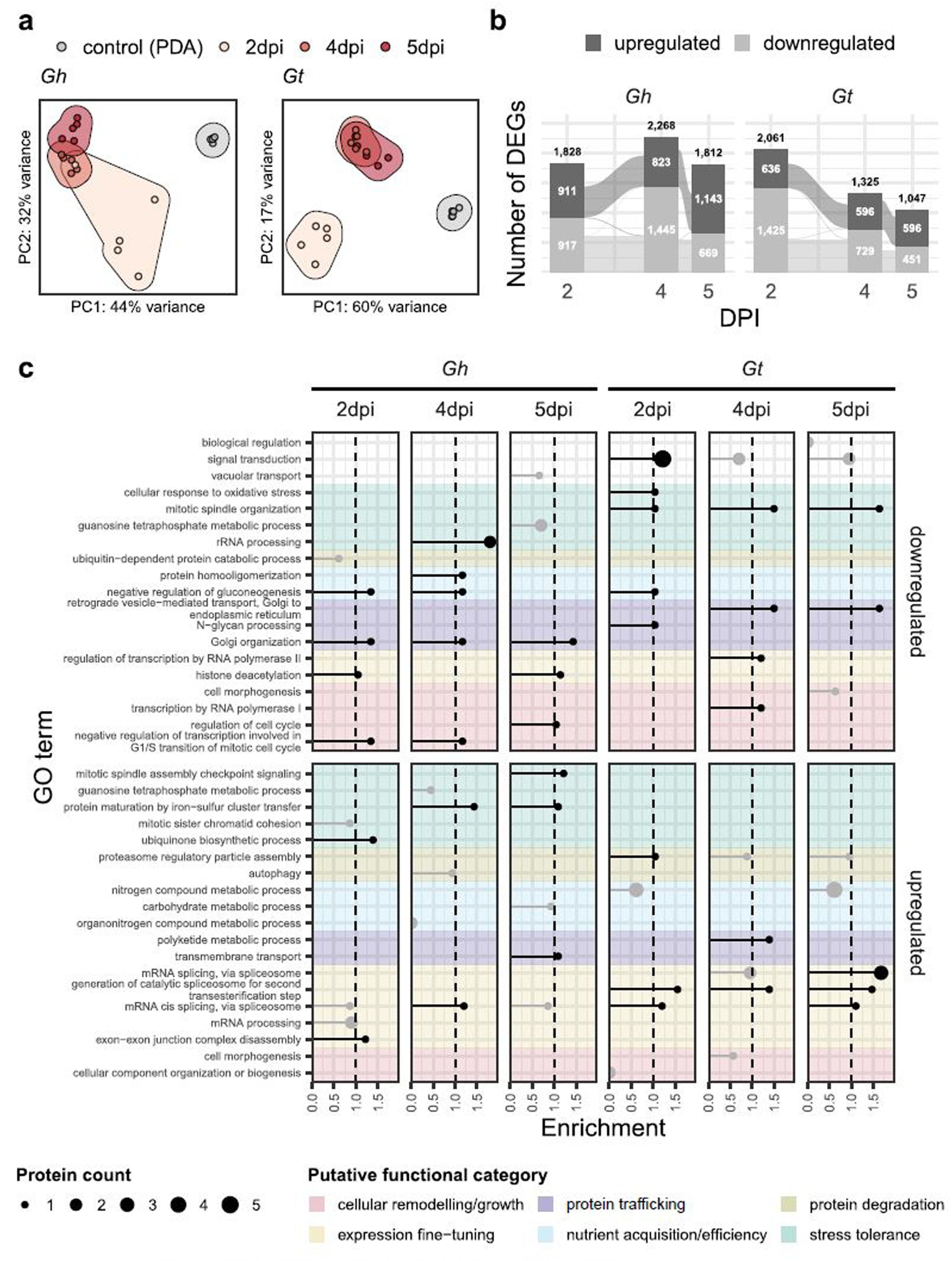
**(a)** Variance-stabilised PCAs showing the expression profiles of the top 500 most variable genes for both *G. tritici* (*Gt*) and *G. hyphopodioides* (*Gh*), coloured according to time point. **(b)** Number of significantly up- and down-regulated genes in each fungus at each time point. Connecting flows between bars indicate where the same genes were differentially expressed across time points. **(c)** Top 5 enriched GO terms for the up- or down-regulated genes in each fungus at each time point. Protein count is represented by the point size and those points crossing the dashed line and in black are deemed statistically significant (-log10(p-value)).

Interestingly, in terms of the overall number of differentially expressed genes (DEGs), *G. tritici* had reduced downregulation as the infection progressed (Fig. 3b). Moreover, the significantly upregulated DEGs at day 4 and day 5 were largely overlapping. However, *G. hyphopodioides* initially increased the overall level of differential expression from 2 dpi to 4 dpi (∼1,000 more DEGs than *G. tritici*) and, despite also reducing the overall number of DEGs from 4 dpi to 5 dpi, there was a considerable increase in upregulated genes at 5 dpi.

We assessed the location of DEGs along the chromosomes, as regional expression patterns have previously been observed in other fungi (e.g. Zhao et al. 2014). Both up- and downregulated genes were broadly scattered across all chromosomes, however, without any evidence of major biases in expression localisation (Supplementary Fig. S4), which is perhaps unsurprising considering previous observations of the ‘one-compartment’ nature of *Gaeumannomyces* genomes (Hill et al. 2025).

### Contrasting transcriptional strategies: *G. hyphopodioides* stress response vs. *G. tritici* expression fine-tuning

We used GO term enrichment analyses to assess the functional roles of DEGs in both the pathogenic *G. tritici* and non-pathogenic *G. hyphopodioides* across the time course. The biological processes associated with significantly up- and down-regulated genes were typically contrasting between species, although processes involving mRNA splicing were upregulated for both (Fig. 3c).

Significantly upregulated GO terms for *G. hyphopodioides* were predominantly associated with stress tolerance, namely: protein maturation by iron-sulfer cluster transfer, ubiquinione biosynthetic process and mitotic spindle assembly checkpoint signalling (Fig. 3c). Other significantly upregulated terms for *G. hyphopodioides* included exon-exon junction complex disassembly at 2 dpi and transmembrane transport at 5 dpi. Significantly downregulated GO terms for *G. hyphopodioides* were frequently the same at multiple time points, including histone deacetylation, negative regulation of gluconeogenesis, Golgi organisation, and negative regulation of G1/S transcription.

Significantly upregulated GO terms for *G. tritici* included: proteasome regulatory particle assembly at 2 dpi and polyketide metabolic process at 4 dpi; and many biological processes involving mRNA splicing at all time points. Significantly downregulated GO terms across time points for *G. tritici* were mostly associated with stress tolerance (mitotic spindle organisation, cellular response to oxidative stress) and protein trafficking (N-glycan processing, retrograde vesicle-mediated transport). A higher number of gene associated with signal transduction were also downregulated at 2 dpi.

### Host interaction factors are broadly upregulated in the non-pathogen

We compared expression profiles of features that are likely to be implicated in the plant-fungal interaction – candidate secreted effector proteins (CSEPs), carbohydrate active enzymes (CAZymes) and putatively characterised biosynthetic gene clusters (BGCs) – between pathogenic *G. tritici* and non-pathogenic *G. hyphopodioides*. CSEPs were predominantly upregulated in *G. hyphopodioides* and only increasing with time, whereas a similar proportion of CSEPs were upregulated and downregulated in *G. tritici*, with the highest overall number at 2 dpi and highly similar CSEP DEGs between 4 dpi and 5 dpi (Fig. 4a). As a set, CSEPs were significantly enriched amongst upregulated genes in *G. hyphopodioides* at all time points, but only at 2 dpi for *G. tritici* (Fig. 4b). Single-copy CSEPs that were shared between *G. hyphopodioides* and *G. tritici* frequently had similar expression patterns in both fungi, but a number that were only upregulated in *G. hyphopodioides* (mostly at 4 and 5 dpi) or downregulated in one fungus but upregulated in the other are interesting contenders for further exploration of what distinguishes lifestyle (Fig. 4e).

**Figure 4.**
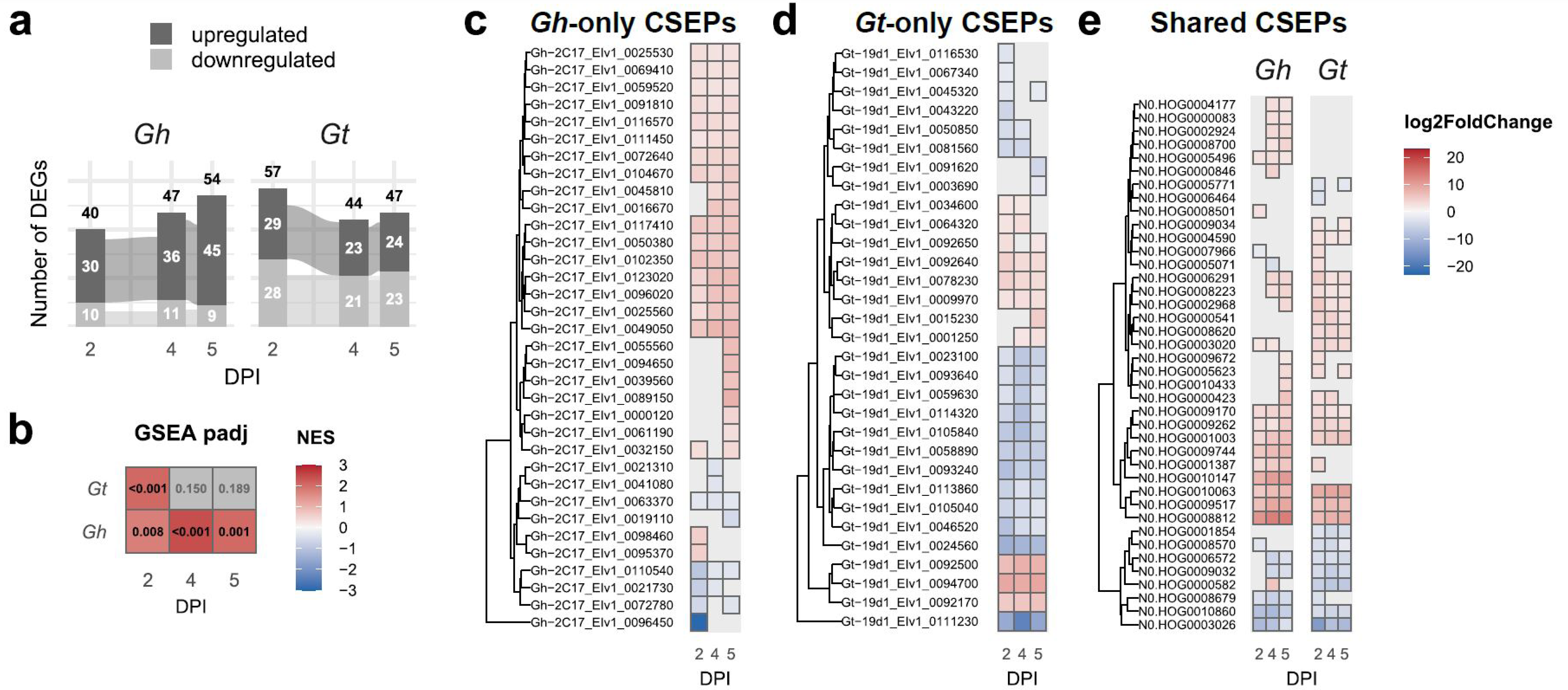
**(a)** Number of significantly up- and down-regulated candidate secreted effector proteins (CSEPs) in each fungus at each time point. *Gh* = *G. hyphopodioides*, *Gt* = *G. tritici*. **(b)** Matrix of adjusted p-values from gene set enrichment analyses (GSEA) showing whether CSEPs as a set are significantly enriched in each fungus at each time point. High normalised enrichment score (NES) indicates enrichment amongst upregulated genes while low NES indicates enrichment amongst downregulated genes. Grey boxes indicate non-significant padj. Expression profiles for differentially expressed CSEPs found only in *G. hyphopodioides* **(c)**, only in *G. tritici* **(d)** and shared in both species (single-copy orthogroups) **(e)**. Genes/orthogroups are ordered by hierarchical clustering, shown to the left of matrices.

There was a similar pattern for CAZymes, with a decrease in DEGs across time points for *G. tritici* and an increase for *G. hyphopodioides*, and a notable jump in upregulated CAZymes at 5 dpi for the latter (Fig. 5a). However, gene set enrichment analysis did not find CAZymes as a set, or those degrading any particular substrate, to be enriched amongst DEGs of either species. There were more species-specific CAZyme DEGs in *G. hyphopodioides* compared to *G. tritici*, predominantly associated with cellulose, polyphenol, xylan and xyloglucan (Fig. 5b). Four out of five lignin-degrading CAZyme DEGs specific to *G. hyphopodioides* were upregulated at 5 dpi (Fig. 5b). The one lignin-degrading DEG that was identified in *G. tritici*, which also belonged to a single-copy orthogroup shared with *G. hyphopodioides*, was upregulated in *G. tritici* and downregulated in *G. hyphopodioides* (Fig. 5d). Three previously characterised lignin-degrading laccases in *G. tritici* (Litvintseva and Henson 2002) were not captured by the CAZyme annotation, but manually checking the homologues of these published genes found that LAC1 was significantly downregulated in both species at 2 dpi and LAC2 was significantly upregulated in *G. hyphopoidoides* at 5 dpi. One broad-spectrum CAZyme orthogroup (N0.HOG0005091), an alpha-d-xyloside xylohydrolase known to act on alpha−galactan, polyphenol, starch, sucrose and xyloglucan, was upregulated at 5 dpi in *G. hyphopodioides* but downregulated at all time points in *G. tritici*.

**Figure 5.**
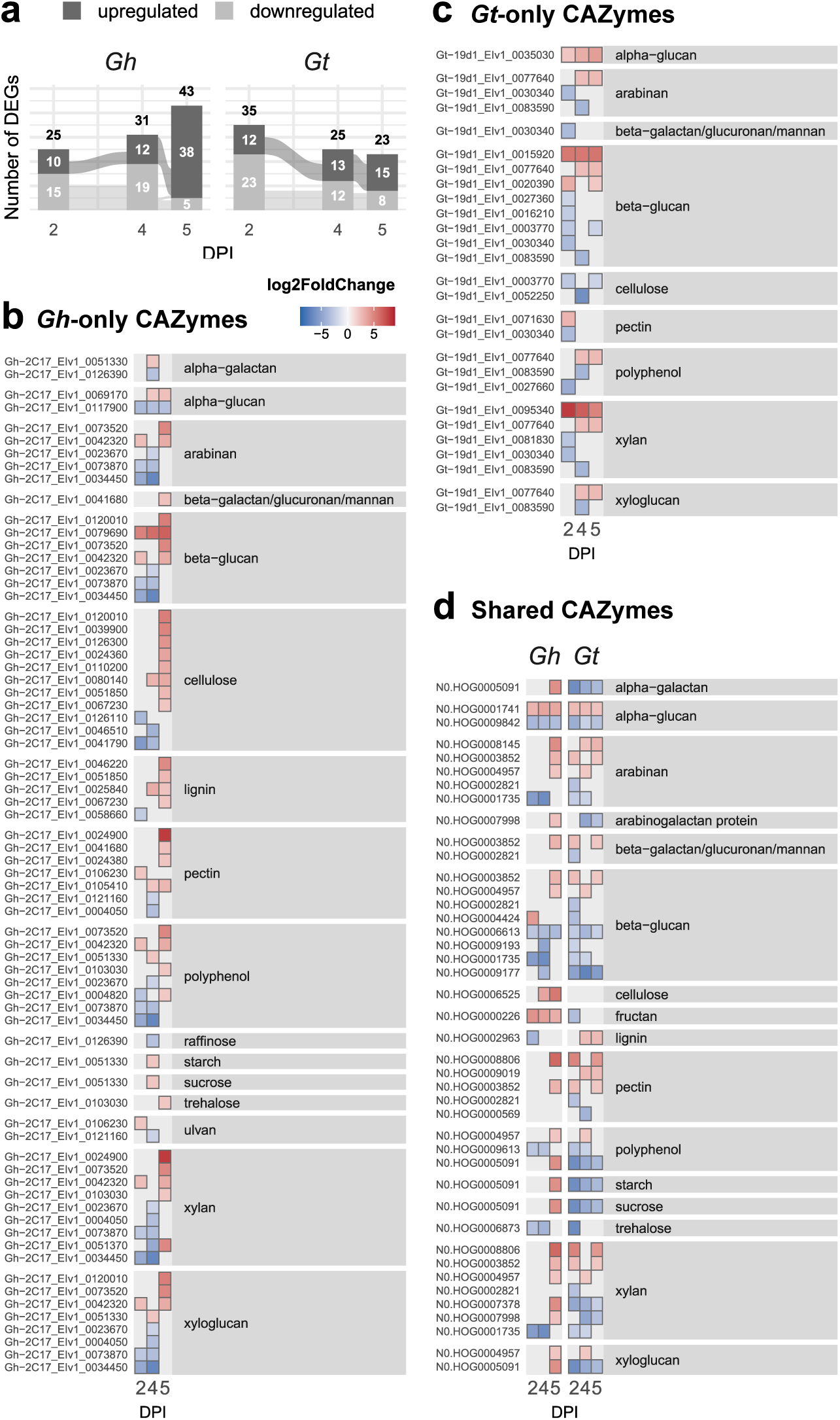
Expression profiles for differentially expressed CAZymes which could be attributed to a substrate, found only in *G. hyphopodioides* **(a)**, only in *G. tritici* **(b)** and shared in both species (single-copy orthogroups) **(c)**. Genes/orthogroups are grouped by the substrate which a CAZyme is known to act on (grey filled boxes). Only plant-derived CAZyme substrates are shown, see Supplementary Fig. S5 for all substrates.

Finally we checked expression profiles of BGCs which have been assigned putative identities based on homology with other species (Hill et al. 2025). Of the two indole BGCs that are unique to *G. hyphopodioides*, one had significant upregulation for three out of five genes, including the key biosynthetic indole gene, across all time points (Fig. 6a). Of the two BGCs that are unique to *G. tritici*, for dichlorodiaporthin twelve out of twenty genes were downregulated at 2 dpi, some of which were also downregulated at 4 dpi, while for equisetin an oxidase, enoylreductase and the biosynthetic PKS-NRPS gene were upregulated across various time points (Fig. 6b). The DHN melanin BGC, which is present in both fungi, had downregulation for five out of six genes in *G. tritici*, including the PKS biosynthetic gene, but no apparent pattern of differential expression for *G. hyphopodioides* (Fig. 6c). The reverse was true of nectriapyrone, which was downregulated for half of the genes including the biosynthetic PKS and an O-methyltransferase in *G. hyphopodioides*, but had no significant DEGs in *G. tritici*. The clavaric acid BGC showed a comparable expression profile of downregulation across the two fungi, although marginally more pronounced for *G. tritici* (Fig. 6c).

**Figure 6.**
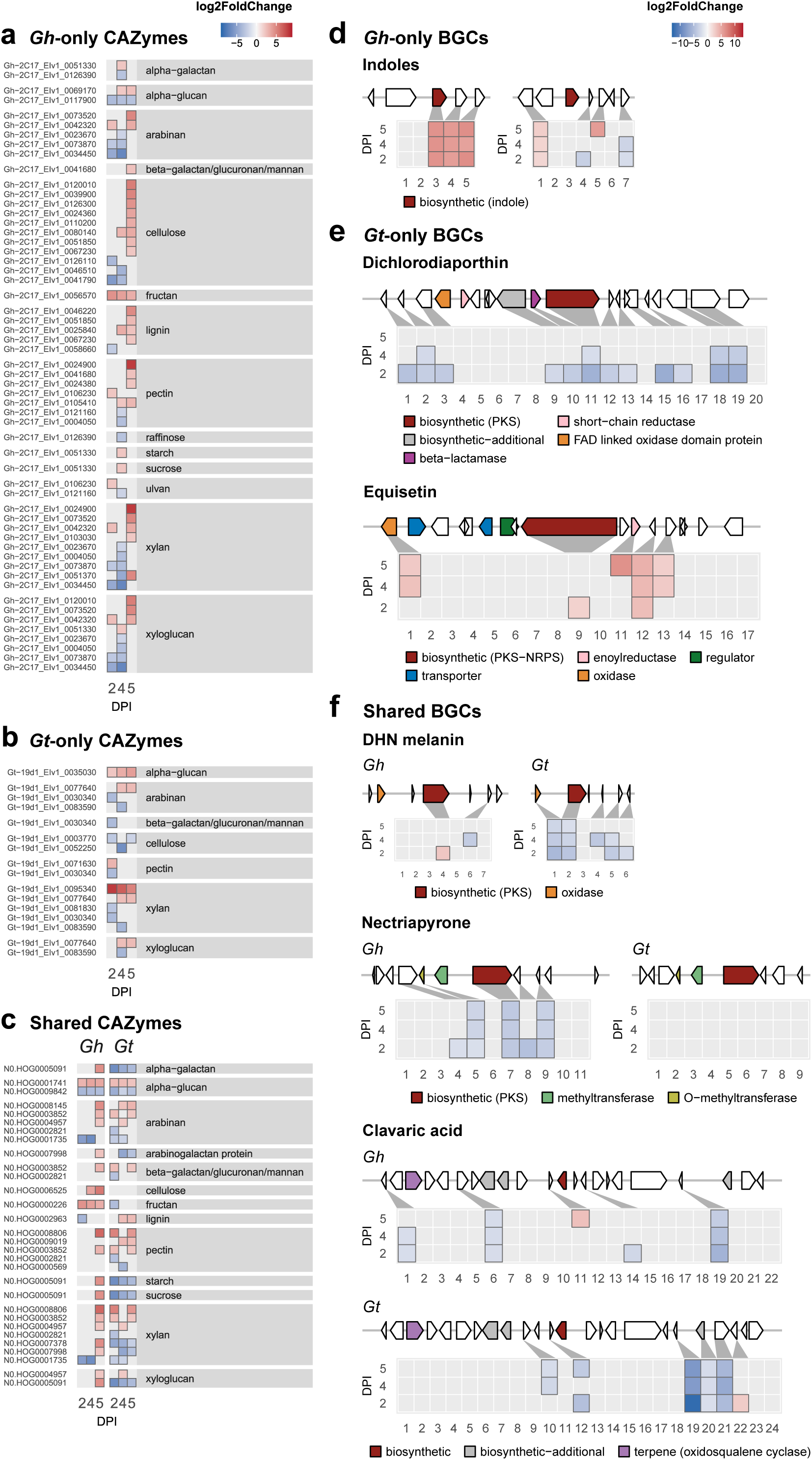
Expression profiles of biosynthetic gene clusters with putative functional annotations (Hill et al. 2025), found only in *G. hyphopodioides* **(d)**, only in *G. tritici* **(e)** and shared in both species **(f)**. For other uncharacterised BGCs see Supplementary Fig. S6.

## Discussion

Wheat take-all, the most important wheat root disease, has been difficult to study at the transcriptional level due to limited fungal genomic resources. By integrating new *G. tritici* and *G. hyphopodioides* reference genomes (Hill et al. 2025) with RNA-seq data from inoculated plants (Chancellor et al. 2024), we assessed the potential of dual host-fungal transcriptomes to compare expression dynamics of two divergent fungal lifestyles during early-stage colonisation. Microbial transcripts are often recovered in low abundance compared to host transcripts in dual RNA-seq datasets (e.g. Dobon et al. 2016; Chung et al. 2021). Despite the anticipated host bias, we found early time points to have a low rate of mis-mapping (i.e. minimal reads from uncolonised wheat samples erroneously mapping to fungal genomes) and the expected distribution of gene expression to be representative of the fungal transcriptome (Supplementary Fig. S2). We recovered an increasing proportion of reads for both *G. tritici* and *G. hyphopodioides* across the time course, consistent with progressive colonisation of host roots. However, *G. hyphopodioides* reads at 5 dpi accounted for nearly a quarter of total reads, six times greater than *G. tritici* at this time point. This relative increase in the number and proportion of *G. hyphopodioides* read data at 5 dpi may reflect increasing fungal biomass as *G. hyphopodioides* forms the dense, swollen SEVs in the cortex. But without later time points we cannot confirm if or when we might see a similar but delayed pattern for *G. tritici*, as can be observed for some pathogens when comparing resistant and susceptible hosts (e.g. Shi et al. 2020).

### Transcriptional patterns are mirrored in host and endophyte and support SEV formation being symptomatic of fungal stress

Chancellor et al. (2024) originally characterised the wheat host transcriptional response to early colonisation by the pathogenic *G. tritici* versus the non-pathogenic fungal endophyte *G. hyphopodioides*. The authors found host interaction with *G. hyphopodioides* to be particularly remarkable as it resulted in local endophyte-mediated resistance (EMR). Extensive transcriptional reprogramming of the wheat host was observed in *G. hyphopodioides*-colonised tissue consistent with that of an immune response, including enrichment of antifungal cinnamic acid, increased response to wounding and regulation of defence responses such as lignin biosynthesis (Chancellor et al. 2024; Speakman and Lewis 1978). The role of lignification in plant immunity is well-established: it provides a physical barrier to slow initial pathogen colonisation, such as the lignitubers which form to enclose invading hyphae, and also owing to the antimicrobial phytoalexins which accumulate along the same pathway (Ninkuu et al. 2022; Miedes et al. 2014).

The dramatic increase in the number of host differentially expressed genes in *G. hyphopodioides*-colonised tissue across 4 and 5 dpi were reflected in our results here from the fungal side showing pronounced upregulation of *G. hyphopodioides* genes occurring at the same time points (Fig. 3b). This was concurrent with SEV formation in the fungus. SEVs have been characterised in numerous *Magnaporthaceae* species and morphological similarities have been drawn between the “microsclerotia” of dark septate endophytes, typically found in stressful conditions (Ban et al. 2012; Knapp et al. 2018), and the chlamydospores of some pathogenic fungi (Chancellor 2022). This suggests potential for SEVs to be long-term survival structures (Francisco et al. 2019) and does not necessarily rule out the potential for opportunistic re-infection. The multi-layered, melanised cell walls and high numbers of putative lipid bodies contained within SEVs may provide *G. hyphopodioides* with a well-protected lipid source to survive periods of stress during host colonisation (Chancellor 2022). Perhaps surprisingly, however, genes in the BGC for DHN melanin were largely not differentially expressed in *G. hyphopodioides* (Fig. 6c), although this may be an artefact of the experimental design: differential expression was inferred relative to the fungus grown in potato dextrose broth, in which it is likely to be already producing melanin (Wang et al. 2020). Any signal of differential expression from SEVs, which are a small subset of the total cells sampled, may also be diluted amongst the bulk RNA-seq.

During the period of *G. hyphopodioides* SEV formation and upregulation of genes (4-5dpi) GO term enrichment also highlighted signals associated with stress in *G. hyphopodioides* (Fig. 3c), likely in response to plant immune factors such as the antifungal precursors, phytoalexin byproducts, and increase in reactive oxygen species associated with lignin biosynthesis (Chancellor et al. 2024; Miedes et al. 2014). Additionally, high levels of host-lignification may restrict *G. hyphopodioides*’ access to carbon sources, which could explain the upregulation (downregulation of negative regulation) of gluconeogenesis, both to compensate for reduced nutrient availability and to support the energetic requirements of SEV formation. This offers evidence in support of the hypothesised model from Chancellor et al. (2024) for SEV production: that induced plant defences cause cessation of hyphal growth, swelling, and melanisation, eventually forming mature SEVs as response to stress and nutrient limitation.

Curiously, SEVs are capable of forming penetration pegs, which allow hyphae to colonise adjacent cells (Chancellor et al. 2024) and has been reported for other dark septate endophytes (Jumpponen and Trappe 1998). Paired with the observation that colonisation by *G. hyphopodioides* following a *G. tritici* invasion was found to worsen the take-all infection (Chancellor et al. 2024), questions are raised about the factors influencing *G. hyphopodioides* as a biocontrol measure. This could be indicative of *Gaeumannomyces* species existing on a pathogenic-mutualistic spectrum, potentially exhibiting lifestyle transience or opportunism, as has been hypothesised for other fungal lineages (Collinge et al. 2022; Hill et al. 2022). Therefore, the notion of biocontrol as a panacea ought to be approached with caution. Further research into the unresolved function of SEVs is required prior to employing *G. hyphopodioides*-mediated resistance as a biocontrol solution to take-all. Regardless of their overall function, we identify that stress signals may contribute to SEV formation, and we establish the synchronicity with which host and fungal expression occur when modulating this endophytic interaction.

### Is G. hyphopodioides in a resting state or actively modulating the plant-fungal interaction?

Perception of the host by fungal colonisers is intimate and complex: it provides key developmental signals but is also inextricable from stress due to both pre-existing and induced host defences (Shalaby and Horwitz 2015). This holds true in our expression analysis of *G. hyphopodioides*, where enrichment of stress responses occurred synchronously with both the host’s spike in gene expression and the formation of SEVs. This, and the aforementioned similarity of SEVs to other structures associated with stress or long-term survival, has pointed towards a hypothesis that SEVs represent a resting state induced by the stress resulting from the localised host immune response (Chancellor et al. 2024).

However, other results here prompt the question as to whether *G. hyphopodioides* is ‘idle’ or engaging more actively with the plant host. For instance, a considerable number of CSEPs and CAZymes are upregulated at 5 dpi, but it is not clear if this is as a result of SEV formation, or in spite of it. The fungus does not progress past the lignified endodermal layer, and yet we see upregulation of lignin-degrading CAZymes, amongst many other plant cell wall degrading enzymes. Another interesting insight was that one of the two indole BGCs present in *G. hyphopodioides* (and absent in *G. tritici*) was consistently upregulated across the time course (Fig. 6a). Fungal indole derivatives are known to act as growth promoters and signalling molecules for multiple plant-fungal interactions, including mutualistic and symbiotic ones (Fu et al. 2015). For instance, *Falciphora oryzae*, a root endophyte in the same family as *Gaeumannomyces* species (*Magnaporthaceae*), upregulates indole derivatives in response to exudates from the host, stimulating *Arabidopsis* lateral root growth (Sun et al. 2020). Furthermore, GO term analysis revealed downregulation of genes which inhibit the transcription of genes involved in the cell cycle, which suggests that DNA replication is being promoted rather than slowed.

While *G. hyphopodioides* is well documented to reliably not cause disease in wheat and provide protection from *G. tritici* infection (e.g. Speakman and Lewis 1978; Martyniuk and Myśków 1984; Osborne et al. 2018), more specific understanding of where the fungus falls on the mutualistic–pathogenic spectrum has not been resolved. For instance, the nutritional strategy of *G. hyphopodioides* is unclear. Are SEVs indeed a resting state enabling the fungus to persist until host senescence, at which point it acts as a saprotroph to decompose plant cells? Does the fungus receive ‘by-product benefits’ (Ruotsalainen et al. 2022) from the host, driving the interaction towards mutualism? The formation of penetration pegs from the SEVs which may reach the plant plasma membrane could instead point towards *G. hyphopodioides* directly gaining nutrients from living host cells but not so aggressively as to kill the host or impact yield (Osborne et al. 2018), in either a parasitic or mutualistic interaction. Further interrogation of the intricacies of this plant-fungal interaction at the cellular level is needed to shed light on these unresolved questions.

### The transcriptionally stable, stealthy, pathogenicity strategy of G. tritici

Contrary to *G. hyphopodioides*, pathogenic *G. tritici* appeared to remain transcriptionally constant across 4 and 5 dpi. This pattern was consistent throughout the results, with plateaued read counts; 4 and 5 dpi occupying a shared region of PCA space; highly similar differentially expressed genes across 4 and 5 dpi; and high overlap in GO term enrichment overlap between the two time points. This stable expression was surprising, as the corresponding confocal microscopy from Chancellor et al. (2024) identified clear colonisation progression across these time points, where *G. tritici* surpassed the endodermal barrier and colonised the root stele. This may suggest that despite penetrating through different root tissue sections, its colonisation strategy remains constant. Another key difference between *G. tritici* and *G. hyphopodioides* is that we do not see evidence of *G. tritici* exhibiting the stress response which was identified in *G. hyphopodioides* even though each enters and colonises the same host and tissue. Rather, for *G. tritici* there was downregulation of stress associated GO terms and DHN melanin, a secondary metabolite associated with stress protection (Henson et al. 1999). Additionally, a greater proportion of *G. tritici* CSEPs were downregulated compared to *G. hyphopodioides*. The expression profile was dominated by downregulation relative to the non-pathogen, suggesting that the early colonisation strategy of *G. tritici* may rely on stealth to minimise host detection. Downregulation of signal transduction genes, the expression of which could trigger host recognition (Buscaill and van der Hoorn 2021), would also support this idea. This hypothesis of early reliance on stealth corresponds with the lack of major host transcriptional reprogramming in the *G. tritici*-colonised samples (Chancellor et al. 2024).

While the transcriptional response of the host to *G. tritici* was not as pronounced as *G. hyphopodioides*, *G. tritici* infection did also cause some degree of host lignification (Chancellor et al. 2024). This does not prevent disease, however, suggesting that the pathogen manages to evade or overcome these local host defences (Chancellor et al. 2024). The lignin-degrading laccase enzymes have also previously been implicated in *G. tritici* pathogenesis and protection from host oxidative defences (Chancellor 2022; Yang et al. 2015b; Daval et al. 2011; Edens et al. 1999). Yang et al. (2015b) found upregulation of two of the three characterised *G. tritici* laccase genes during wheat root infection, and downregulation of the third, however here we only found one of these laccase genes to be significantly differentially expressed, which was downregulated at 2 dpi. We did, however, identify a differentially expressed laccase orthogroup shared between *G. tritici* and *G. hyphopodioides* which is apparently novel as it was not homologous with any of the previously mentioned *G. tritici* laccase genes. This laccase was upregulated in *G. tritici* at 4 and 5 dpi when the fungus successfully penetrates the stele and represented the only case of a shared CAZyme which was upregulated in *G. tritici* but downregulated in *G. hyphopodioides* (Fig. 5d), making it a key target for further study of lifestyle determinants between these species.

Polyketide metabolic process was another significantly upregulated GO term for *G. tritici*, and polyketides are known to include many pathogenic toxins and virulence factors (Chooi and Tang 2012). The equisetin BGC, which is present in *G. tritici* but not in *G. hyphopodioides*, was found to have a number of genes significantly upregulated, and is known in two *Fusarium* pathosystems to cause necrotic lesions of roots and other tissues of crops including cereals (Wheeler et al. 1999), suggesting it may be another key virulence factor in take-all. Perhaps counterintuitively, DHN melanin genes were downregulated in *G. tritici*, despite melanin being previously implicated in its pathogenicity on wheat (Freeman and Ward 2004; Henson et al. 1999; Aranda et al. 2023). As there is an energetic cost to producing secondary metabolites such as melanin, there is a trade-off between its production and the fungus’ growth rate (Siletti et al. 2017), and at this key early stage *G. tritici* may favour rapid growth to ‘outrun’ plant defences and invade the stele. Incidentally, the non-pathogenic *G. hyphopodioides* is known to be more heavily melanised than *G. tritici* (Henson et al. 1999), and the fact that melanin plays a role in both stress protection as well as a structural or mechanical role indicates that the function of melanin in the colonisation/infection dynamics of these fungi is more nuanced.

### Future directions

While this dataset captures the early divergence in expression during initial colonisation of the pathogenic *G. tritici* and non-pathogenic *G. hyphopodioides* in wheat, it does not capture the progression of the interaction post day 5. The time points in this dataset are key to differentiating the role of SEVs and their importance in pathogenicity, or lack thereof, but a longer time course would allow more insight into the nuances of maintaining these contrasting lifestyles. Moreover, it would be particularly interesting to determine whether *G. tritici* undergoes a transcriptional shift, akin to *G. hyphopodioides*, at a later time point, for instance when the fungus shifts from stealth to active necrosis, producing lesions. Using fixed time points (days post inoculation) for different fungi, or even replicates for a single fungus, whose colonisation rate may not be strictly circadian, may be over-simplistic or even misleading. This is supported by the fact that one of our 2 dpi replicates for *G. hyphopodioides* was closer to the 4 dpi replicates in PCA space, suggesting that the colonisation for that sample had progressed more rapidly. This scenario could be explored for by comparing expression trajectories, instead of fixed points, over a longer period.

As previously discussed, low fungal read count at early stages is a common obstacle for early infection multi-species transcriptomic studies (Chung et al. 2021). Emerging single-cell RNA-seq technologies may offer an opportunity to combat this bias at early stages of colonisation where fungal biomass is low, whilst resolving tissue-specific expression of both fungus and host simultaneously. Single-cell and/or spatial RNA-seq would also provide more targeted expression profiles of key regions or structures for colonisation, such as SEVs. Future studies should also consider the medium used for fungal controls, to make the null and *in planta* environments as comparable as possible and minimise the risk of artefacts in the differential expression results.

## Conclusions

This study represents the first use of dual host-fungal RNA-seq to investigate the transcriptional behaviour of both *G. tritici* and *G. hyphopodioides* during early wheat root colonisation. By integrating newly available reference genomes (Hill et al. 2025) with dual host-fungus transcriptomic data, molecular evidence for diverging colonisation strategies between the pathogen *G. tritici* and endophyte *G. hyphopodioides* were revealed. *G. tritici* evaded the host with little response and maintained a largely stable transcriptional profile, progressively reducing downregulation, with potential enzymatic mechanisms for immune evasion. *G. hyphopodioides* halted progression in the cortex and displayed a dynamic transcriptional response between 4 and 5 dpi, mirroring host gene expression and indicative of stress-associated SEV formation. At this stage it is not possible to attribute the pausing in the cortex to either host or endophyte but SEVs remain a key stage to explore in order to understand the causes of cessation of invasion. These results emphasise the importance of exploring non-pathogenic relatives to identify novel insights into the mechanisms underpinning pathogenicity. Comparative transcriptomics of the host, pathogen, and endophyte in this system provide a unique opportunity to explore fungal virulence, stress tolerance, and biocontrol potential for this important crop disease.

## Supporting information

Supplementary Datasheet

## Acknowledgements

We thank Dan Smith, Gail Canning and Javier Palma-Guerrero for their contributions to prior analysis of the plant host data in the dual RNA-seq experiment, and Erika Kroll for assistance depositing the new control data. Thanks also go to Amy Dodd for the illustrations in Figure 1.

## Funding

The authors acknowledge funding from the Biotechnology and Biological Sciences Research Council (BBSRC), part of UK Research and Innovation, Core Capability Grants BB/CCG1720/1 and the work delivered via the Scientific Computing group, as well as support for the physical HPC infrastructure and data centre delivered via the NBI Computing infrastructure for Science (CiS) group. This research was supported by the BBSRC grant Delivering Sustainable Wheat (BB/X011003/1) and the constituent work packages (BBS/E/ER/230003B Earlham Institute and BBS/E/RH/230001B Rothamsted Research). TC was supported by the BBSRC funded University of Nottingham Doctoral Training Programme (BB/M008770/1).

## Notes

### Competing Interest Statement

The authors have declared no competing interest.

## References

Alexa, A., and Rahnenfuhrer, J. 2022.topGO: Enrichment Analysis for Gene Ontology.

Andrews, S. 2018. FastQC: a quality control tool for high throughput sequence data. Available at: http://www.bioinformatics.babraham.ac.uk/projects/fastqc/.

Aranda, C., Méndez, I., Barra, P. J., Hernández-Montiel, L., Fallard, A., Tortella, G., Briones, E., and Durán, P. 2023. Melanin Induction Restores the Pathogenicity of *Gaeumannomyces graminis* var. *tritici* in Wheat Plants. JoF. 9:350 10.3390/jof9030350.

Ban, Y., Tang, M., Chen, H., Xu, Z., Zhang, H., and Yang, Y. 2012. The Response of Dark Septate Endophytes (DSE) to Heavy Metals in Pure Culture. PLoS ONE. 7:e47968 10.1371/journal.pone.0047968.

Benjamini, Y., and Hochberg, Y. 1995. Controlling The False Discovery Rate - A Practical And Powerful Approach To Multiple Testing Article. Journal of the Royal Statistical Society. 57:289–300 10.1111/j.2517-6161.1995.tb02031.x.

Berendsen, R. L., Pieterse, C. M. J., and Bakker, P. A. H. M. 2012. The rhizosphere microbiome and plant health. Trends in Plant Science. 17:478–486 10.1016/j.tplants.2012.04.001.

Buscaill, P., and van der Hoorn, R. A. L. 2021. Defeated by the nines: nine extracellular strategies to avoid microbe-associated molecular patterns recognition in plants. The Plant Cell. 33:2116–2130 10.1093/plcell/koab109.

Chancellor, T. 2022. Evaluating the potential of non-pathogenic Magnaporthaceae species for the control of take-all disease in wheat.

Chancellor, T., Smith, D. P., Chen, W., Clark, S. J., Venter, E., Halsey, K., Carrera, E., McMillan, V., Canning, G., Armer, V. J., Hammond-Kosack, K. E., and Palma-Guerrero, J. 2024. A fungal endophyte induces local cell wall–mediated resistance in wheat roots against take-all disease. Front Plant Sci. 15:1444271 10.3389/fpls.2024.1444271.

Chooi, Y.-H., and Tang, Y. 2012. Navigating the Fungal Polyketide Chemical Space: From Genes to Molecules. J. Org. Chem. 77:9933–9953 10.1021/jo301592k.

Chung, M., Bruno, V. M., Rasko, D. A., Cuomo, C. A., Muñoz, J. F., Livny, J., Shetty, A. C., Mahurkar, A., and Dunning Hotopp, J. C. 2021. Best practices on the differential expression analysis of multi-species RNA-seq. Genome Biol. 22:121 10.1186/s13059-021-02337-8.

Collinge, D. B., Jensen, B., and Jørgensen, H. J. 2022. Fungal endophytes in plants and their relationship to plant disease. Current Opinion in Microbiology. 69:102177 10.1016/j.mib.2022.102177.

Danecek, P., Bonfield, J. K., Liddle, J., Marshall, J., Ohan, V., Pollard, M. O., Whitwham, A., Keane, T., McCarthy, S. A., Davies, R. M., and Li, H. 2021. Twelve years of SAMtools and BCFtools. GigaScience. 10:giab008 10.1093/gigascience/giab008.

Daval, S., Lebreton, L., Gazengel, K., Boutin, M., Guillerm-Erckelboudt, A., and Sarniguet, A. 2011. The biocontrol bacterium *Pseudomonas fluorescens* Pf29Arp strain affects the pathogenesis-related gene expression of the take-all fungus *Gaeumannomyces graminis* var. *tritici* on wheat roots. Molecular Plant Pathology. 12:839–854 10.1111/j.1364-3703.2011.00715.x.

Dobon, A., Bunting, D. C. E., Cabrera-Quio, L. E., Uauy, C., and Saunders, D. G. O. 2016. The host-pathogen interaction between wheat and yellow rust induces temporally coordinated waves of gene expression. BMC Genomics. 17:380 10.1186/s12864-016-2684-4.

Edens, W. A., Goins, T. Q., Dooley, D., and Henson, J. M. 1999. Purification and Characterization of a Secreted Laccase of *Gaeumannomyces graminis* var. *tritici*. Appl Environ Microbiol. 65:3071–3074 10.1128/AEM.65.7.3071-3074.1999.

Fones, H. N., Bebber, D. P., Chaloner, T. M., Kay, W. T., Steinberg, G., and Gurr, S. J. 2020. Threats to global food security from emerging fungal and oomycete crop pathogens. Nature Food. 1:332–342 10.1038/s43016-020-0075-0.

Francisco, C. S., Ma, X., Zwyssig, M. M., McDonald, B. A., and Palma-Guerrero, J. 2019. Morphological changes in response to environmental stresses in the fungal plant pathogen *Zymoseptoria tritici*. Sci Rep. 9:9642 10.1038/s41598-019-45994-3.

Freeman, J., and Ward, E. 2004. *Gaeumannomyces graminis*, the take-all fungus and its relatives. Mol Plant Pathol. 5:235–252 10.1111/j.1364-3703.2004.00226.x.

Freeman, J., Ward, E., Gutteridge, R. J., and Bateman, G. L. 2005. Methods for studying population structure, including sensitivity to the fungicide silthiofam, of the cereal take-all fungus, Gaeumannomyces graminis var. tritici. Plant Pathol. 54:686–698 10.1111/j.1365-3059.2005.01252.x.

Fu, S.-F., Wei, J.-Y., Chen, H.-W., Liu, Y.-Y., Lu, H.-Y., and Chou, J.-Y. 2015. Indole-3-acetic acid: A widespread physiological code in interactions of fungi with other organisms. Plant Signaling & Behavior. 10:e1048052 10.1080/15592324.2015.1048052.

Henson, J. M., Butler, M. J., and Day, A. W. 1999. The Dark Side of the Mycelium: Melanins of Phytopathogenic Fungi. Annu. Rev. Phytopathol. 37:447–471 10.1146/annurev.phyto.37.1.447.

Hill, R., Buggs, R. J. A., Vu, D. T., and Gaya, E. 2022. Lifestyle Transitions in Fusarioid Fungi are Frequent and Lack Clear Genomic Signatures. Mol Biol Evol. 39:msac085 10.1093/molbev/msac085.

Hill, R., Grey, M., Fedi, M. O., Smith, D., Canning, G., Ward, S. J., Irish, N., Smith, J., McMillan, V. E., Hammond, J., Osborne, S.-J., Reynolds, G., Smith, E., Chancellor, T., Swarbreck, D., Hall, N., Palma-Guerrero, J., Hammond-Kosack, K. E., and McMullan, M. 2025. Evolutionary genomics reveals variation in structure and genetic content implicated in virulence and lifestyle in the genus *Gaeumannomyces*. BMC Genomics. 26:239 10.1186/s12864-025-11432-0.

Jumpponen, A., and Trappe, J. M. 1998. Dark septate endophytes: a review of facultative biotrophic root-colonizing fungi. New Phytologist. 140:295–310 10.1046/j.1469-8137.1998.00265.x.

Kim, D., Paggi, J. M., Park, C., Bennett, C., and Salzberg, S. L. 2019. Graph-based genome alignment and genotyping with HISAT2 and HISAT-genotype. Nat Biotechnol. 37:907–915 10.1038/s41587-019-0201-4.

Knapp, D. G., Németh, J. B., Barry, K., Hainaut, M., Henrissat, B., Johnson, J., Kuo, A., Lim, J. H. P., Lipzen, A., Nolan, M., Ohm, R. A., Tamás, L., Grigoriev, I. V., Spatafora, J. W., Nagy, L. G., and Kovács, G. M. 2018. Comparative genomics provides insights into the lifestyle and reveals functional heterogeneity of dark septate endophytic fungi. Scientific Reports. 8:6321 10.1038/s41598-018-24686-4.

Korotkevich, G., Sukhov, V., Budin, N., Shpak, B., Artyomov, M. N., and Sergushichev, A. 2016. Fast gene set enrichment analysis. 10.1101/060012.

Kovaka, S., Zimin, A. V., Pertea, G. M., Razaghi, R., Salzberg, S. L., and Pertea, M. 2019. Transcriptome assembly from long-read RNA-seq alignments with StringTie2. Genome Biol. 20:278 10.1186/s13059-019-1910-1.

Kwak, Y.-S., and Weller, D. M. 2013. Take-all of Wheat and Natural Disease Suppression: A Review. Plant Pathol J. 29:125–135 10.5423/PPJ.SI.07.2012.0112.

Liao, Y., and Shi, W. 2020. Read trimming is not required for mapping and quantification of RNA-seq reads at the gene level. NAR Genomics and Bioinformatics. 2:lqaa068 10.1093/nargab/lqaa068.

Litvintseva, A. P., and Henson, J. M. 2002. Cloning, Characterization, and Transcription of Three Laccase Genes from *Gaeumannomyces graminis* var. *tritici*, the Take-All Fungus. Appl Environ Microbiol. 68:1305–1311 10.1128/AEM.68.3.1305-1311.2002.

Love, M. I., Huber, W., and Anders, S. 2014. Moderated estimation of fold change and dispersion for RNA-seq data with DESeq2. Genome Biol. 15:550 10.1186/s13059-014-0550-8.

Malicka, M., Magurno, F., and Piotrowska-Seget, Z. 2022. Plant association with dark septate endophytes: When the going gets tough (and stressful), the tough fungi get going. Chemosphere. 302:134830 10.1016/j.chemosphere.2022.134830.

Martyniuk, S., and Myśków, W. 1984. Control of the Take-all Fungus by *Phialophora* sp. (lobed hyphopodia) in Microplots Experiments with Wheat. Zentralblatt für Mikrobiologie. 139:575–579 10.1016/S0232-4393(84)80075-X.

Mc Carthy, U., Uysal, I., Badia-Melis, R., Mercier, S., O’Donnell, C., and Ktenioudaki, A. 2018. Global food security – Issues, challenges and technological solutions. Trends in Food Science & Technology. 77:11–20 10.1016/j.tifs.2018.05.002.

Mehrabi, Z., McMillan, V. E., Clark, I. M., Canning, G., Hammond-Kosack, K. E., Preston, G., Hirsch, P. R., and Mauchline, T. H. 2016. *Pseudomonas* spp. diversity is negatively associated with suppression of the wheat take-all pathogen. Sci Rep. 6:29905 10.1038/srep29905.

Miedes, E., Vanholme, R., Boerjan, W., and Molina, A. 2014. The role of the secondary cell wall in plant resistance to pathogens. Front. Plant Sci. 5:358 10.3389/fpls.2014.00358.

Ninkuu, V., Yan, J., Fu, Z., Yang, T., Ziemah, J., Ullrich, M. S., Kuhnert, N., and Zeng, H. 2022. Lignin and Its Pathway-Associated Phytoalexins Modulate Plant Defense against Fungi. JoF. 9:52 10.3390/jof9010052.

Osborne, S.-J., McMillan, V. E., White, R., and Hammond-Kosack, K. E. 2018. Elite UK winter wheat cultivars differ in their ability to support the colonization of beneficial root-infecting fungi. J Exp Bot. 69:3103–3115 10.1093/jxb/ery136.

Palma-Guerrero, J., Chancellor, T., Spong, J., Canning, G., Hammond, J., McMillan, V. E., and Hammond-Kosack, K. E. 2021. Take-All Disease: New Insights into an Important Wheat Root Pathogen. Trends Plant Sci. 26:836–848 10.1016/j.tplants.2021.02.009.

R Core Team. 2024.R: A language and environment for statistical computing.

Raaijmakers, J. M., Paulitz, T. C., Steinberg, C., Alabouvette, C., and Moënne-Loccoz, Y. 2009. The rhizosphere: a playground and battlefield for soilborne pathogens and beneficial microorganisms. Plant Soil. 321:341–361 10.1007/s11104-008-9568-6.

Ruotsalainen, A. L., Kauppinen, M., Wäli, P. R., Saikkonen, K., Helander, M., and Tuomi, J. 2022. Dark septate endophytes: mutualism from by-products? Trends in Plant Science. 27:247–254 10.1016/j.tplants.2021.10.001.

Savary, S., Willocquet, L., Pethybridge, S. J., Esker, P., McRoberts, N., and Nelson, A. 2019. The global burden of pathogens and pests on major food crops. Nat Ecol Evol. 3:430–439 10.1038/s41559-018-0793-y.

Shalaby, S., and Horwitz, B. A. 2015. Plant phenolic compounds and oxidative stress: integrated signals in fungal–plant interactions. Curr Genet. 61:347–357 10.1007/s00294-014-0458-6.

Shi, W., Zhao, S.-L., Liu, K., Sun, Y.-B., Ni, Z.-B., Zhang, G.-Y., Tang, H.-S., Zhu, J.-W., Wan, B.-J., Sun, H.-Q., Dai, J.-Y., Sun, M.-F., Yan, G.-H., Wang, A.-M., and Zhu, G.-Y. 2020. Comparison of leaf transcriptome in response to *Rhizoctonia solani* infection between resistant and susceptible rice cultivars. BMC Genomics. 21:245 10.1186/s12864-020-6645-6.

Siletti, C. E., Zeiner, C. A., and Bhatnagar, J. M. 2017. Distributions of fungal melanin across species and soils. Soil Biology and Biochemistry. 113:285–293 10.1016/j.soilbio.2017.05.030.

Speakman, J. B., and Lewis, B. G. 1978. Limitation of *Gaeumannomyces graminis* by wheat root responses to *Phialophora radicicola*. New Phytologist. 80:373–380 10.1111/j.1469-8137.1978.tb01571.x.

Sun, X., Wang, N., Li, P., Jiang, Z., Liu, X., Wang, M., Su, Z., Zhang, C., Lin, F., and Liang, Y. 2020. Endophytic fungus *Falciphora oryzae* promotes lateral root growth by producing indole derivatives after sensing plant signals. Plant Cell Environ. 43:358–373 10.1111/pce.13667.

Wang, M., Ren, X., Wang, L., Lu, X., Han, L., Zhang, X., and Feng, J. 2020. A functional analysis of mitochondrial respiratory chain cytochrome *bc*_1_ complex in *Gaeumannomyces tritici* by RNA silencing as a possible target of carabrone. Molecular Plant Pathology. 21:1529–1544 10.1111/mpp.12993.

Weller, D. M., Raaijmakers, J. M., McSpadden Gardener, B. B., and Thomashow, L. S. 2002. Microbial populations responsible for specific soil suppressiveness to plant pathogens. Annu Rev Phytopathol. 40:309–348 10.1146/annurev.phyto.40.030402.110010.

Wheeler, M. H., Stipanovic, R. D., and Puckhaber, L. S. 1999. Phytotoxicity of equisetin and epi-equisetin isolated from *Fusarium equiseti* and *F. pallidoroseum*. Mycol Res. 103:967–973 10.1017/S0953756298008119.

Wickham, H. 2016.ggplot2: Elegant Graphics for Data Analysis.

Williams, C. R., Baccarella, A., Parrish, J. Z., and Kim, C. C. 2016. Trimming of sequence reads alters RNA-Seq gene expression estimates. BMC Bioinformatics. 17:103 10.1186/s12859-016-0956-2.

Yang, L., Han, X., Zhang, F., Goodwin, P. H., Yang, Y., Li, J., Xia, M., Sun, R., Jia, B., Zhang, J., Quan, X., Wu, C., Xue, B., and Lu, C. 2018. Screening *Bacillus* species as biological control agents of *Gaeumannomyces graminis* var. *Tritici* on wheat. Biological Control. 118:1–9 10.1016/j.biocontrol.2017.11.004.

Yang, L., Quan, X., Xue, B., Goodwin, P. H., Lu, S., Wang, J., Du, W., and Wu, C. 2015a. Isolation and identification of *Bacillus subtilis* strain YB-05 and its antifungal substances showing antagonism against *Gaeumannomyces graminis* var. *tritici*. Biological Control. 85:52–58 10.1016/j.biocontrol.2014.12.010.

Yang, L., Xie, L., Xue, B., Goodwin, P. H., Quan, X., Zheng, C., Liu, T., Lei, Z., Yang, X., Chao, Y., and Wu, C. 2015b. Comparative Transcriptome Profiling of the Early Infection of Wheat Roots by *Gaeumannomyces graminis* var. *tritici* D. Zhou, ed. PLoS ONE. 10:e0120691 10.1371/journal.pone.0120691.

Yun, Y., Yu, F., Wang, N., Chen, H., Yin, Y., and Ma, Z. 2012. Sensitivity to silthiofam, tebuconazole and difenoconazole of *Gaeumannomyces graminis* var. *tritici* isolates from China. Pest Management Science. 68:1156–1163 10.1002/ps.3277.

Zhao, C., Waalwijk, C., de Wit, P. J., Tang, D., and van der Lee, T. 2014. Relocation of genes generates non-conserved chromosomal segments in *Fusarium graminearum* that show distinct and co-regulated gene expression patterns. BMC Genomics. 15:191 10.1186/1471-2164-15-191.

Zhu, T., Wang, L., Rimbert, H., Rodriguez, J. C., Deal, K. R., De Oliveira, R., Choulet, F., Keeble-Gagnère, G., Tibbits, J., Rogers, J., Eversole, K., Appels, R., Gu, Y. Q., Mascher, M., Dvorak, J., and Luo, M. 2021. Optical maps refine the bread wheat *Triticum aestivum* cv. Chinese Spring genome assembly. The Plant Journal. 107:303–314 10.1111/tpj.15289.

