## Supplementary Material for "Transcriptional signatures underlying divergent lifestyles of endophytic and pathogenic fungi in early colonisation of wheat roots"

Stella Moren-Rosado[1], Rowena Hill[1], Tania Chancellor[2], Rachel Rusholme Pilcher[1], Neil Hall[1,3], Kim Hammond-Kosack[4], and Mark McMullan[1]

[1]*Earlham Institute, Norwich Research Park, Colney Ln, Norwich, NR4 7UZ, UK*
[2]*Royal Botanic Gardens, Kew, Richmond, Surrey, TW9 3AB, UK*
[2]*School of Biological Sciences, University of East Anglia, Norwich, NR4 7TJ, UK*
[4]*Harnessing Biotic Interactions, Rothamsted Research, Harpenden, Herts, Al5 2JQ, UK*

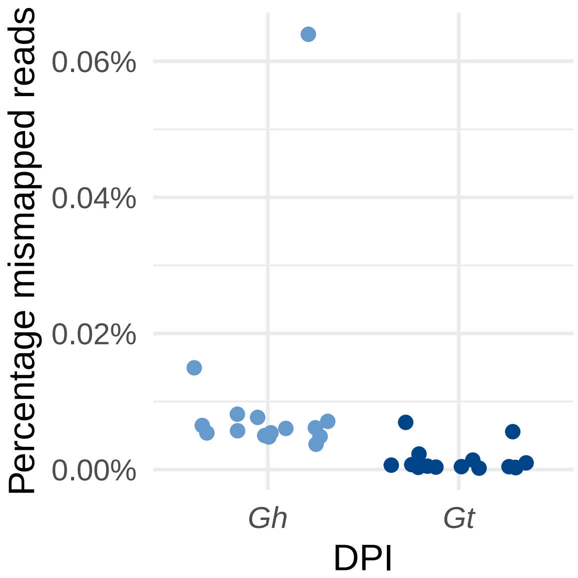


**Supplementary Fig. S1** Proportion of the total RNA reads of control samples (i.e. no fungal inoculation) which mapped to the fungal reference genomes.

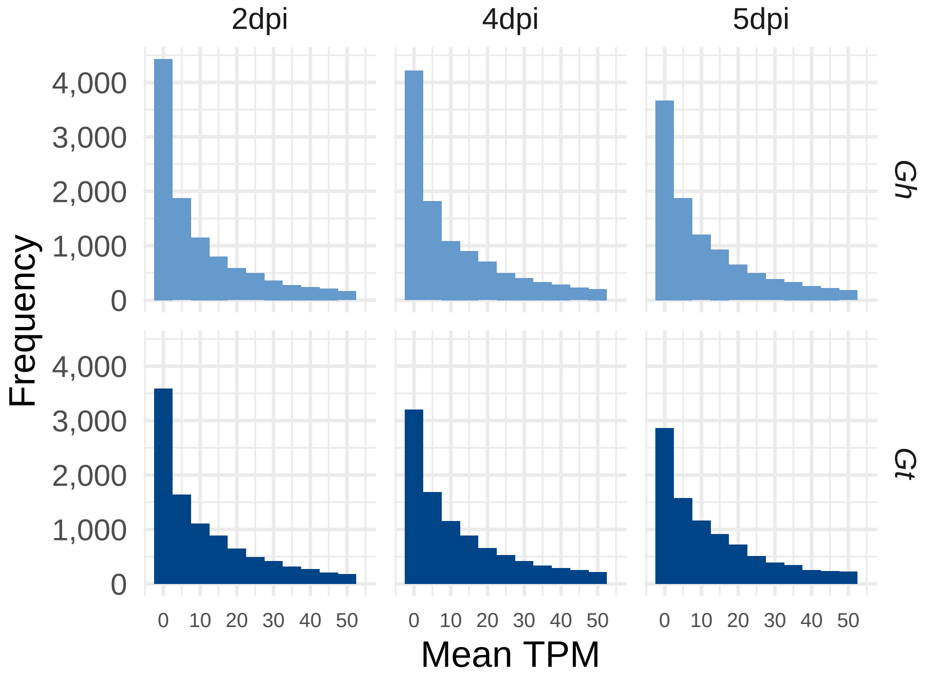


**Supplementary Fig. S2** Histograms showing the average transcripts per million (TPM) for each gene at each time point.

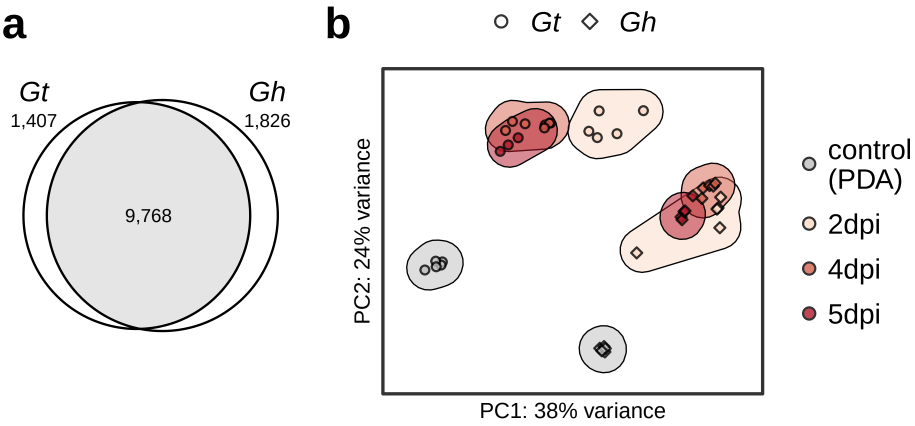


**Supplementary Fig. S3 (a)** Venn diagram showing the common single-copy orthogroups shared by *Gt* and *Gh*. **(b)** Variance-stabilised PCA showing the expression profiles of the top 500 most variable shared single-copy orthogroups for both *Gt* and *Gh* at each to time point

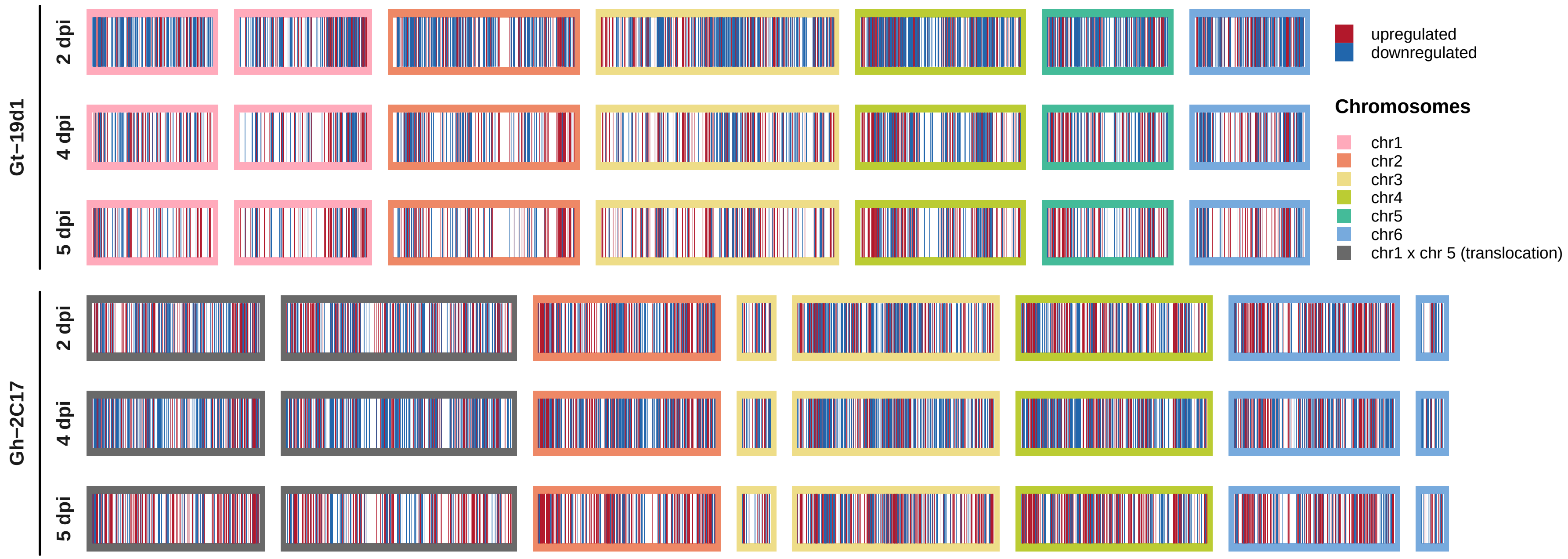


**Supplementary Fig. S4** Location of differentially expressed genes across chromosomes.

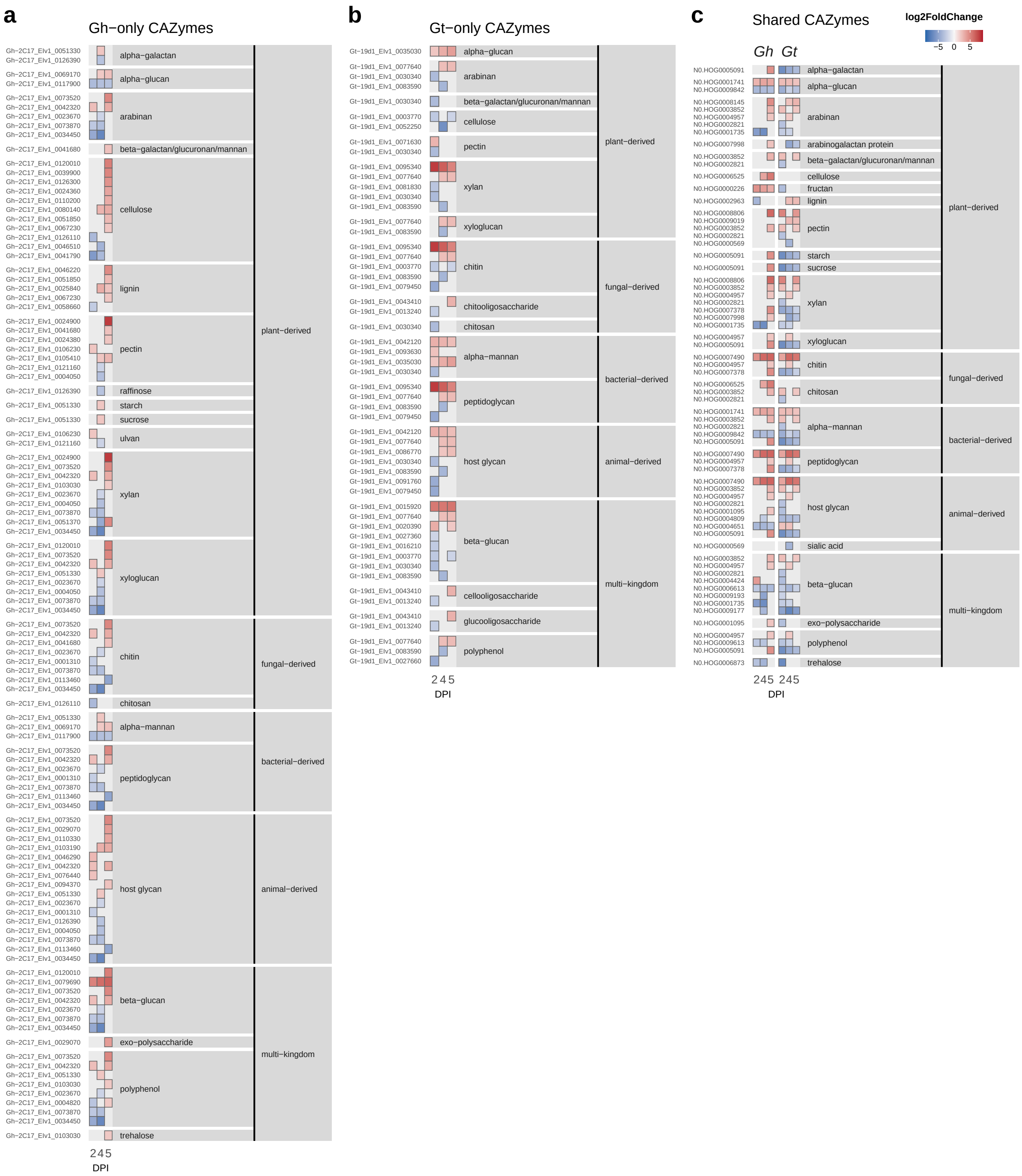


**Supplementary Fig. S5** Expression profiles for differentially expressed CAZyme genes which could be attributed to a substrate, found only in *Gh* **(a)**, only in *Gt* **(b)** and shared in both species (single-copy orthogroups) **(c)**.

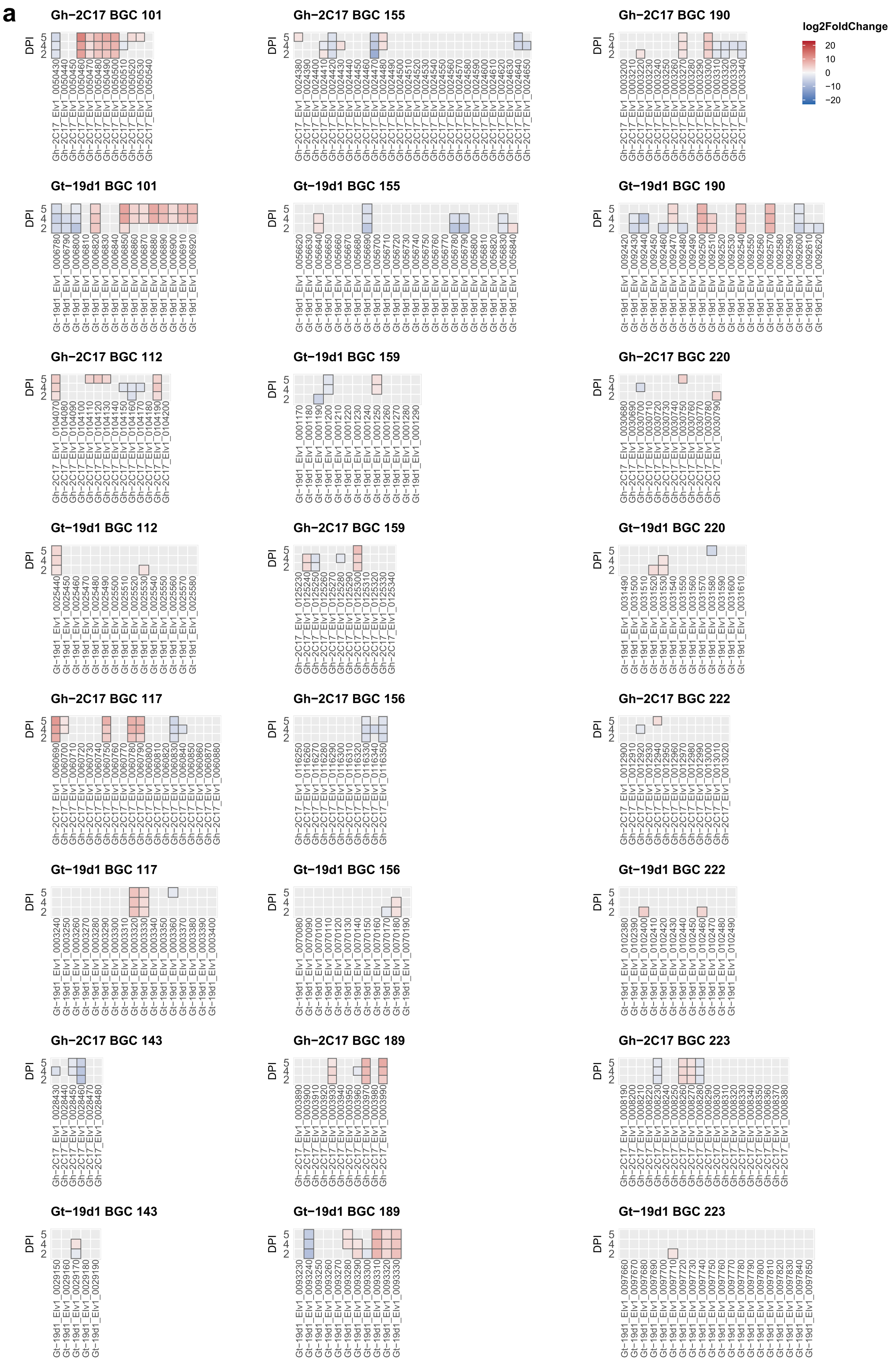


**Supplementary Fig. S6** Expression profiles for uncharacterised biosynthetic gene clusters (BGCs), shared in both species **(a)**, found only in *Gt* **(b)**, and found only in *Gh* **(c)**. ▼

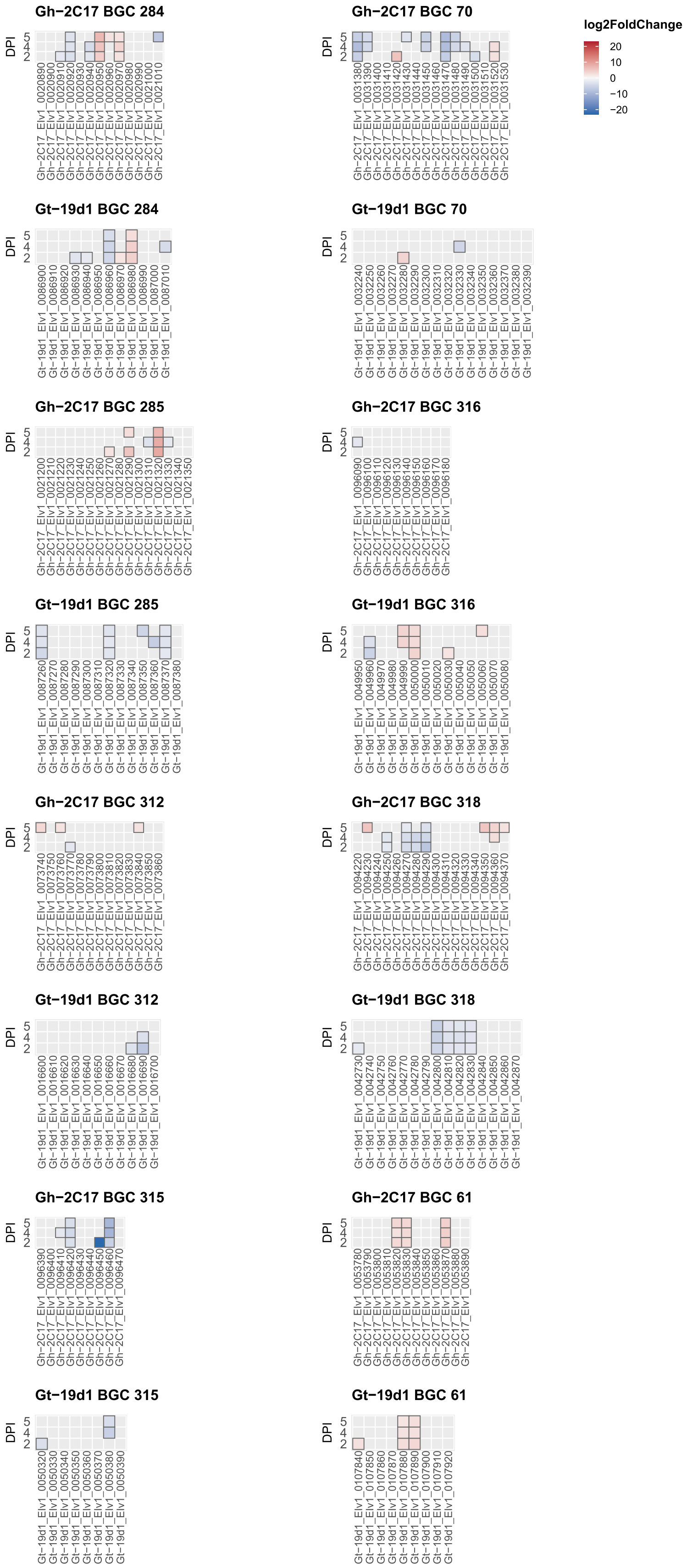


**Supplementary Fig. S6 continued. ▼**

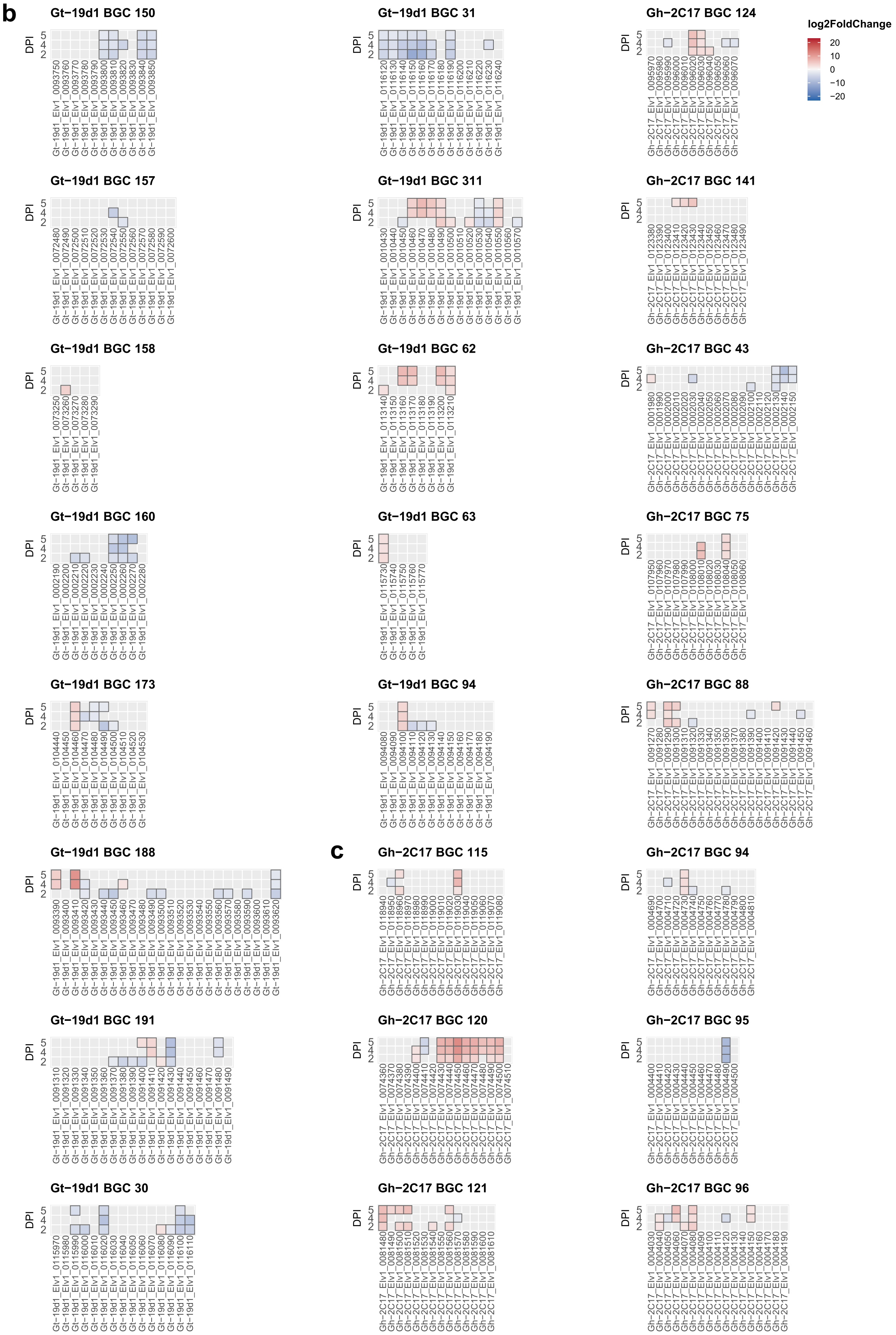


**Supplementary Fig. S6 continued.**
